# Early EEG and behavioral alterations in Dravet mice

**DOI:** 10.1101/2020.01.13.904557

**Authors:** Saja Fadila, Shir Quinn, Ana Turchetti Maia, Daniel Yakubovich, Karen L. Anderson, Moran Rubinstein

## Abstract

Dravet Syndrome (Dravet) is a severe childhood epileptic encephalopathy. The disease begins around the age of six months, with a febrile stage, characterized by febrile seizures with otherwise normal development. By the end of the first year of life, the disease progresses to the worsening stage, featuring recurrent intractable seizures and the appearance of additional comorbidities, including global developmental delay, cognitive deficits, hyperactivity and motor problems. Later, in early school years, Dravet reaches the stabilization stage, in which seizure burden decreases, while Dravet-associated comorbidities persist. Dravet syndrome mouse models (DS) faithfully recapitulate the three stages of the human syndrome. Here, we performed power spectral analyses of background EEG activity in DS and their wild-type (WT) littermates, demonstrating disease stage-related alterations. Specifically, while the febrile stage activity resembled that of WT mice, we observed a marked reduction in total power during the worsening stage and a smaller reduction during the stabilization stage. Moreover, low EEG power at the worsening stage correlated with increased risk for premature death, suggesting that such measurements can potentially be used as a marker for Dravet severity. With normal development at the febrile stage and the presentation of developmental delay at the worsening stage, the contribution of recurrent seizures to the emergence of Dravet-associated comorbidities is still debated. Thus, we further characterized the behavior of WT and DS mice during the different stages of Dravet. At the febrile stage, despite their normal background EEG patterns, DS mice already demonstrated motor impairment and hyperactivity in the open field, that persisted to the worsening and stabilization stages. Conversely, clear evidence for deficits in working memory emerged later in life, during the worsening stage. These results indicate that despite the mild epilepsy at the febrile stage, DS development is already altered, suggesting that the pathophysiological mechanisms governing the appearance of some Dravet behavioral comorbidities may be independent of the epileptic phenotype.

**Highlights:**

- Reduction in background EEG power in Dravet
- Low EEG power correlates with the risk of premature death
- Motor deficits and hyperactivity are evident as early as the febrile stage
- Cognitive deficits and detection of increased anxiety begin at the worsening stage

## Introduction

Dravet Syndrome (Dravet) is a rare, severe childhood-onset epileptic encephalopathy, caused by *de novo* mutations in the *SCN1A* gene (Claes et al., 2002). Dravet patients develop normally in their first months of life, but subsequently exhibit unusually severe febrile seizures. Towards the end of the first year, the epilepsy progresses to severe refractory spontaneous seizures and frequent episodes of status epilepticus. Clinical improvement of the seizure phenotype is usually observed after six years of age, with a reduction in the frequency of spontaneous seizures. Based on the severity of epileptic phenotype, the disease can be divided into ‘febrile’, ‘worsening’ and ‘stabilization’ stages (Gataullina and Dulac, 2017; Wirrell et al., 2017). Importantly, in addition to severe intractable seizures, the worsening stage is characterized by the presentation of additional comorbidities, including global developmental delay, moderate-to-severe intellectual disability, deficits in perception, memory, verbal abilities, motor development and ataxia (Dravet and Oguni, 2013).

The causal link between the epileptic phenotype and associated comorbidities in Dravet is still debated. On the one hand, development is normal during the febrile stage and deteriorates later during the worsening stage (Gataullina and Dulac, 2017; Wirrell et al., 2017). Moreover, early onset of spontaneous seizure and status epilepticus, before the end of the first year of life, were correlated with a worse developmental outcome (Brunklaus et al., 2012; Cetica et al., 2017). Conversely, several prospective clinical studies identified early developmental alterations, demonstrating deficits in visuomotor function before the onset of spontaneous seizures (Battaglia et al., 2016; Chieffo et al., 2011; Guzzetta, 2011), as well as delayed mastery of independent sitting (Verheyen et al., 2019), indicating early development is not completely normal. Furthermore, other studies have shown dissociation between the severity of epilepsy and cognitive abilities (Nabbout et al., 2013; Ouss et al., 2019).

To date, there are seven different mouse models for Dravet, based on either truncation or missense mutations in the *Scn1a* gene (Cheah et al., 2012; Miller et al., 2014; Ogiwara et al., 2013, 2007; Ricobaraza et al., 2019; Tsai et al., 2015; Yu et al., 2006). Regardless of the underlying genetic perturbation, all models demonstrate severe epilepsy and premature death. Moreover, adult DS mice were shown to exhibit cognitive deficits, hyperactivity and motor impairment (Han et al., 2012; Ito et al., 2013; Kalume et al., 2007; Ricobaraza et al., 2019; Rubinstein et al., 2015a; Williams et al., 2019), confirming that DS mice faithfully recapitulate the human syndrome.

Here, we examined background EEG activity and the behavioral features of wild-type (WT) and DS mice at time points corresponding to the febrile, worsening and stabilization stages of Dravet. Our data reveal that while normal background EEG patterns are observed at the febrile stage, a marked reduction in power spectral density (PSD) appears in the worsening stage in DS mice. Intriguingly, lower PSD correlated with increased risk for premature death, indicating a possible link between EEG power and the severity of epilepsy. In addition, our behavioral data demonstrated motor deficits and hyperactivity already at the febrile stage, prior to the onset of severe epilepsy, suggesting that pathophysiological mechanisms governing the emergence of Dravet behavioral comorbidities may be dissociated from the epileptic phenotype.

## Methods

### Animals

All animal experiments were approved by the Institutional Animal Care and Use Committee (IACUC) of Tel Aviv University. Mice used in this study were housed in a standard animal facility at the Goldschleger Eye Institute at a constant (22°C) temperature, on 12-hour light/ dark cycles, with *ad libitum* access to food and water.

We used DS mice harboring the global *Scn1a*^A1783V^ mutation. DS mice were generated by crossing conditional *Scn1a*^A1783V^ males (The Jackson Laboratory; Bar Harbor, ME, USA, stock #026133) with CMV-Cre females (stock #006054). We used this breeding scheme due to the location of Cre on the X chromosome (Schwenk et al., 1995). Both male and female offspring were used. Mice that were positive for the flox *Scn1a*^A1783V^ allele, but negative for Cre expression, were not used.

### Surgery

P16 - P49 mice underwent surgery under ketamine / xylazine (191 / 4.25 mg/kg) anesthesia. Carprofen (5 mg/kg) was injected just prior to the procedure and 24 h post-surgery. Postnatal day (P)16 mice were manually fed until P18. Electrode implantation was done as previously described (Kalume et al., 2015; Rubinstein et al., 2015a, 2015b). Briefly, a midline incision was made above the skull and fine silver wire electrodes (130 μm diameter bare; 180 μm diameter coated) were placed at visually identified locations, bilaterally above the somatosensory cortex; a reference electrode was placed on the cerebellum; and a grounding electrode was placed subcutaneously behind the neck, towards the left shoulder. The electrodes were connected to a micro-connector system, and their impedance was 11.8 ± 0.7 kΩ. After electrode placement, the connector was secured with dental cement and the skin closed with sutures. Mice were allowed at least two nights to recover prior to recording. P18 mice were used to examine EEG activity at the end of the febrile stage; ages P21 - P27 (referred to as P21) were used to examine the worsening stage; and mice between P32 - P49 (referred to as P35) were used for the stabilization stage. The data presented in Figs. 1 and 2 were recorded 2 - 4 days post-surgery, with different groups of mice in each stage of the disease. Fig. 3 displays data collected from the same mice at P21 (the worsening stage of Dravet) and again at P35 (the stabilization stage). Of note, the impedance of the electrode in these recordings was similar at both time points (14.1 ± 1.6 kΩ at P21 and 11 ± 1.3 kΩ at P35, *p* = 0.12).

**Figure 1:**
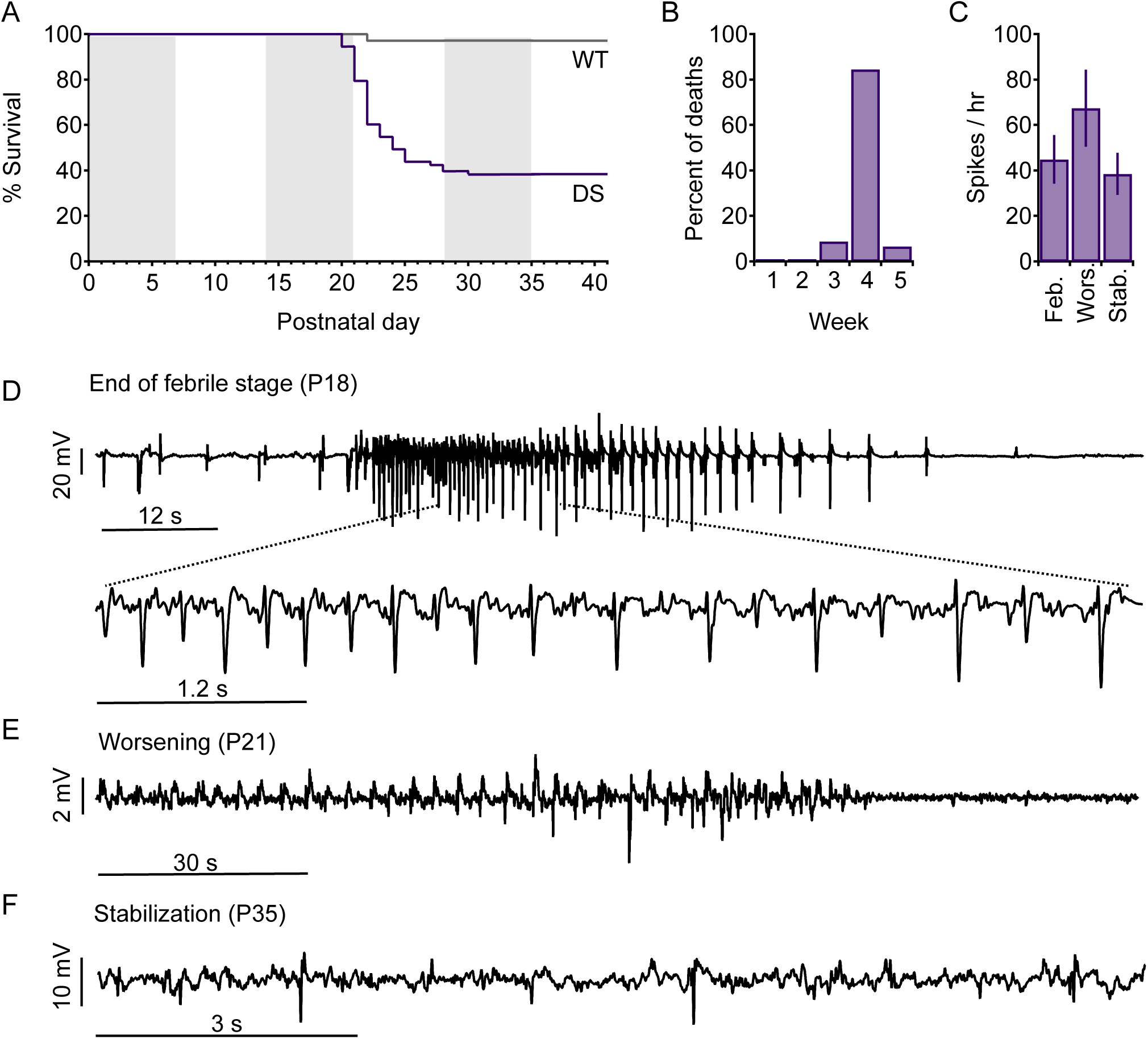
Premature death and epileptic phenotype during Dravet stages. A, Survival plot of DS mice harboring the *Scn1a*^A1783V^ mutation and their WT littermates. WT, n = 71; DS, n = 74. B, Distribution of premature deaths in DS mice. C, Quantification of spikes detected during video - EEG recording of DS mice at the indicated stages. P18, febrile stage; P21 - P27, worsening stage; and P32 - P49, stabilization stage. Spikes were not detected in recordings of WT mice. D, spontaneous seizure in a DS mouse at P18. E, Terminal seizure in P21 DS mouse. F, Epileptic activity in a mouse during the stabilization stage.

**Figure 2:**
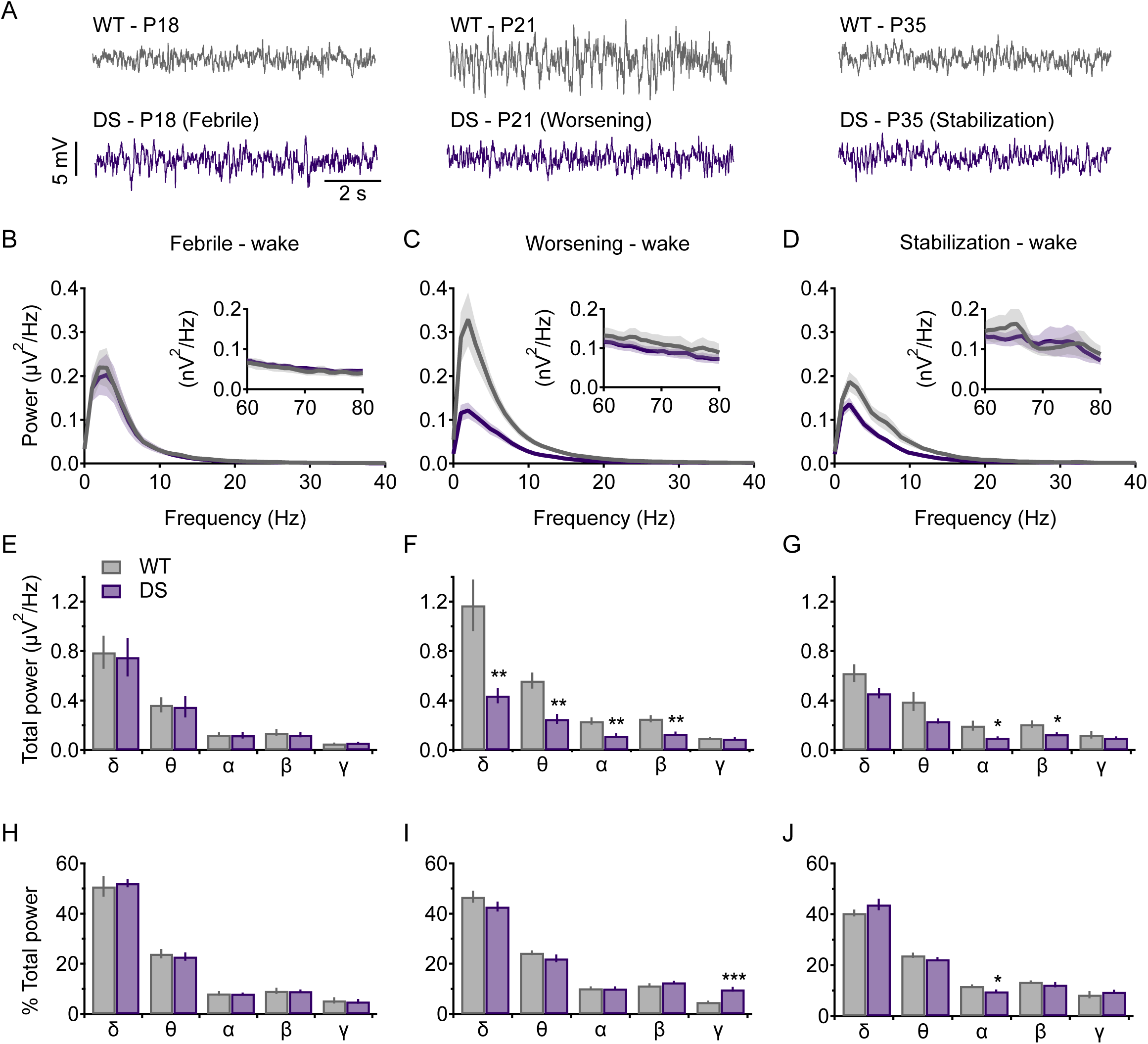
PSD analyses of video - EEG recordings in DS mice. A, Example of WT and DS mouse background EEG traces at the indicated time points. B - D, EEG power density profile during the (B) febrile stage, (C) worsening stage, and (D) stabilization stage. High frequencies (60 - 80 Hz) are separately depicted (insets). E - G. Total power in each frequency band during the (E) febrile, (F) worsening, and (G) stabilization stages. δ: 0.5 - 4 Hz; θ: 4 - 8 Hz; α: 8 - 12 Hz; β: 12 - 30 Hz; γ: 30 - 100 Hz. H -J, Relative power in each frequency band at the (H) febrile, (I) worsening and (J) stabilization stages. Febrile stage (P18): WT: n = 6; DS: n = 6; worsening stage (P21 - P27): WT: n = 13; DS; n = 13, stabilization stage (P32 - P49): WT, n = 12, DS, n = 11. \**p* < 0.05, \*\**p* < 0.01.

**Figure 3:**
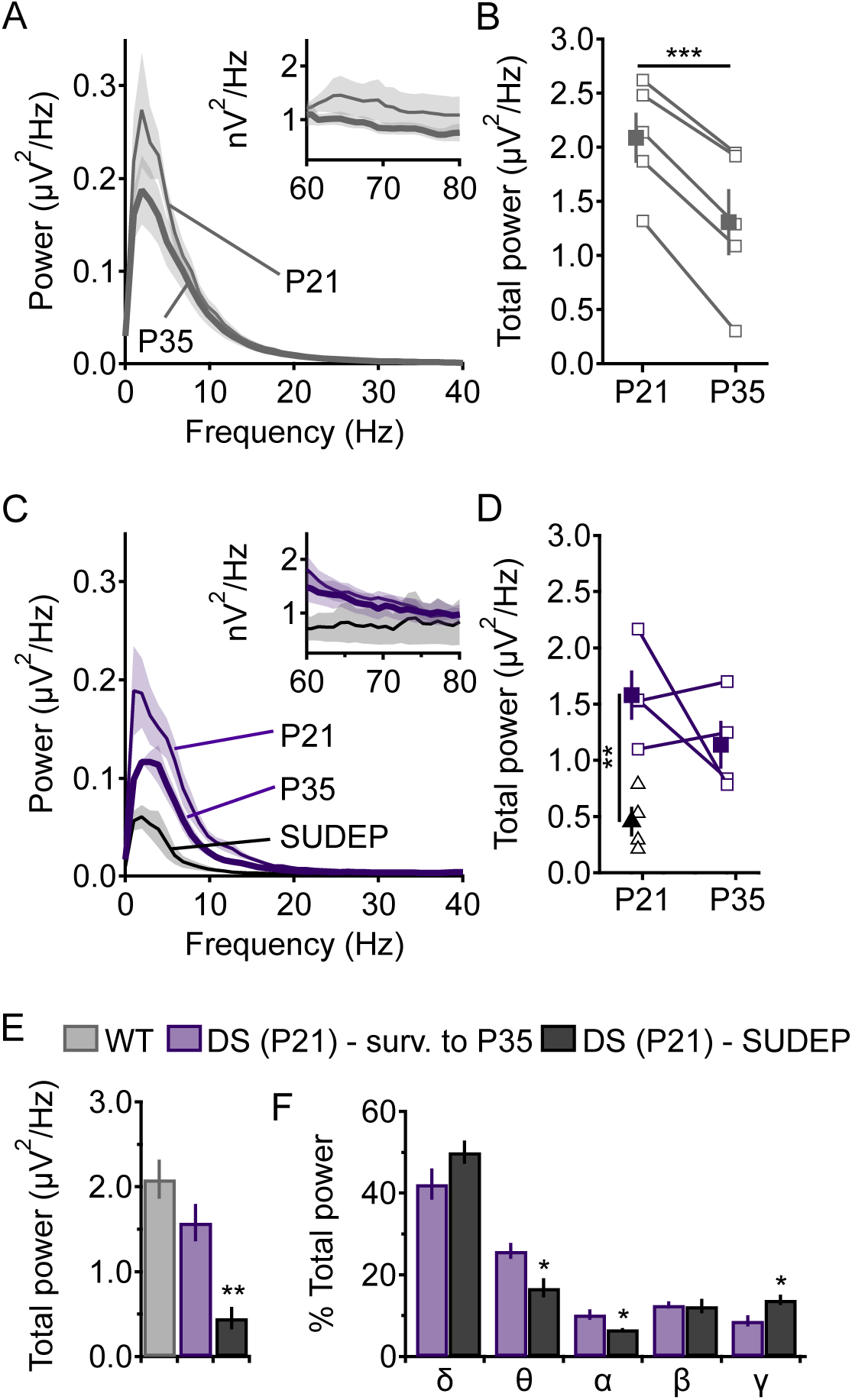
Developmental changes in EEG power. A, EEG power density profile of WT mice at P21 and P35. High frequencies (60 - 80 Hz) are separately depicted (inset). B, Alteration in the total EEG power (0 - 100 Hz) over time in individual WT mice. Empty symbols represent data of individual mice; filled symbols depict average and SE. C, EEG power density profile of DS mice recorded during the worsening (P21) and stabilization (P35) stages, as well as EEG recordings from DS mice that later died prematurely during the worsening stage. High frequency data are presented as in (A). D, Alteration in total EEG power over time. Empty squares represent data of individual mice; empty triangles depict mice that subsequently died during the worsening stage; filled symbols depict average and SE. E, Average total power, at the time point that corresponds to the worsening stage, in WT and DS as indicated. F, Comparison of relative power in each frequency band at the worsening stage between DS mice that survived to the stabilization stage and DS mice that died prematurely. WT, n = 5; DS that survived to the stabilization stage, n = 4, DS mice that died prematurely, n = 4. \**p* < 0.05, \*\**p* < 0.01, \*\*\**p* < 0.001.

### Video - EEG recording and analysis

Video - EEG recordings lasting 3 - 8 hours were obtained from freely behaving mice, connected to a T8 Headstage (Triangle BioSystems, Durham, NC, USA), using a PowerLab 8/35 acquisition hardware and the LabChart 8 software (ADInstruments, Sydney, Australia). The electrical signals were recorded and digitized at a sampling rate of 1 KHz with a notch filter at 50 Hz. The analysis was performed using LabChart 8 (ADInstruments, Sydney, Australia). EEG signals were processed offline with a 100 Hz lowpass filter. Power spectral density was calculated using fast Fourier transform with Hann (cosine-bell) data window set to 50% overlap. The data presented in Figs. 2 and 3 focus on the wake state. To that end, we selected EEG segments that followed animal movement (as seen on video). For each mouse, five to ten 30 s segments devoid of movement artifact were analyzed and averaged. These sections were separated by at least 10 min, and were selected from throughout the entire recording. Spikes were detected using LabChart 8 software (ADInstruments, Sydney, Australia). The threshold was set to 7 standard divisions, and their maximal allowed duration was set to 250 ms.

### Early motor activity

WT and DS mice were tested from P8 to P11. Mice were individually placed in the middle of a 13 cm wide circle and the latency to cross the drawn borders of the circle was measured. The test was repeated three times and averaged. If the mouse was unable to cross the border of the circle within 1 min, the test was terminated.

### Negative geotaxis

Mice (P8 and P11) were placed on an inclined surface (45°), with their head facing downwards. The time to turn and face upward was recorded. If the mouse fell to the bottom of the surface, or did not turn within 1 min, the test was terminated.

### Rotarod

The Rotarod (San Diego Instruments, San Diego, CA, USA) was used to test motor-coordination. Mice (P14 - P35) were placed on an accelerating rotating rod (acceleration from 3 to 40 RPM over 3 min), and the time at which each mouse fell was recorded. The test was repeated five times for each mouse, and the three longest trials were averaged.

### Home cage voluntary wheel running

Mice were individually housed in cages equipped with a Low-Profile Wireless Running Wheel (Med Associates, Fairfax, VT, USA) paired to a hub (DIG-804 USB interface hub). The sum of wheel spins was collected by SOF-860-wheel manager software (Med Associates, Fairfax, VT, USA).

### Open-field test

Mice were placed in the center of a square (50 x 50 cm) Plexiglas apparatus. The open field apparatus was cleaned with 70% ethanol, then water, and wiped dry with paper towels between each trial. Rodent movement was recorded for 10 min by a USB webcam (LifeCam HD-6000, Microsoft, Redmond, WA, USA) and analyzed offline using a video tracking software (EthoVision XT 13, Noldus Technology, Wageningen, Netherlands). For analysis of anxiety-like behavior, the software was set to subdivide the open field into 10 x 10 cm squares (25 in total). Those squares immediately adjacent to the apparatus walls constituted an “outer” zone, while the remaining squares that did not touch the walls demarcated an “inner” zone. DS mice that exhibited seizure activity during or just prior to testing were excluded from analysis.

### Y-Maze spontaneous alteration test

The symmetrical Y-maze comprised three opaque white Plexiglas arms (each 35 cm L x 7.6 cm W x 20 cm H), and was cleaned with 70% ethanol and water, then wiped dry in-between mice. For the test, a mouse was placed into one of the Y-maze arms and allowed free exploration for 10 min. Live tracking was achieved via a monochrome camera (Basler acA1300-60gm, Basler AG, Ahren, Germany) connected with EthoVision XT 13 software (Noldus Technology, Wageningen, Netherlands). The initial placement into a maze arm (designated A, B or C) rotated with each mouse so as to avoid introducing location bias. The analysis includes the number of arms entered, excluding the first recorded arm into which the mouse was placed. The sequence of entries was recorded, as well as the number of triads (entries into a set of 3 different arms, in any sequence). The percentage spontaneous alternation was calculated as the number of triads divided by the number of possible triads.

### Novel object recognition test

The novel object recognition (NOR) test was conducted to assess visuospatial memory. Live tracking of WT and DS littermates utilized a monochrome camera (Basler acA1300-60gm, Basler AG, Ahrensburg, Germany) connected to EthoVision XT 13 software that provided 3-point detection (center-point, nose-point and tail-base). The arena consisted of a round bucket (36 cm floor diameter). To eliminate olfactory cues, the apparatus and the objects were cleaned with 70% ethanol and water, then wiped dry after each trial. Mice were habituated to the test apparatus over 5 days as follows: mice were handled 5 minutes each for 2 consecutive days. On the third and fourth days, mice were allowed to habituate to the bucket for 2 hrs, as a group. On the fifth day, each mouse was individually habituated to the empty bucket for 10 min. Testing occurred on the sixth day, between P21 - P24 and comprised 3 stages. A mouse was first placed into the empty bucket and allowed to explore for 5 min. Thereafter the mouse was returned to its home cage. The second stage took place shortly afterward, after two identical objects (orange bottle caps, 4.8 cm diameter) had been attached to the bucket floor. The test mouse was then placed into the center of the apparatus and allowed to freely explore for 10 min, after which it was returned to its home cage. The third stage occurred shortly thereafter: a third bottle cap (identical to the prior 2) was swapped in as a ‘familiar’ object, and a square white-yellow-black Lego block composite (3.3 cm wide x 3.3 cm high) with smooth sides and a textured upper surface replaced one of the bottle caps as a designated ‘novel’ object. The test mouse was returned to the center of the bucket and allowed to explore freely for 10 min. Exploration of an object was counted when the animal’s nose was located within the area of the object or its surrounding zone (2 cm surrounding the object). Mice that spent less than 20 s exploring both objects were excluded from analysis. If a mouse sat immobile for 10 sec or more, the trial was concluded at that time. Novelty preference was calculated as: Total time (sec) spent exploring the novel object / (Total exploration time of familiar and novel objects) × 100.

### Statistics

Data are reported as mean ± SE. WT and DS mice were compared using unpaired two-tailed Student’s t-test, whereas comparisons of the same mice, recorded at different ages (Fig. 3) utilized a paired two-tailed Student’s t-test. Analysis of variance (ANOVA) was used to examine the statistical significance in Fig. 3E. Differences were considered significant at *p* < 0.05.

## Results

### DS mice recapitulate the three stages of Dravet

DS mice, harboring the *Scn1a*^A1783V^ missense mutation (Kuo et al., 2019; Ricobaraza et al., 2019; Styr et al., 2019), were asymptomatic for their first weeks of life. Spontaneous seizures were not observed prior to P16 (during routine handling) and were rare prior to P19. At P20, premature death had begun, resulting in ∼ 60% overall mortality of DS mice (Fig. 1A). The majority of deaths occurred between P21 and P27 (84.5%); mice that survived beyond P30 were less likely to die prematurely (Fig. 1A, B). Thus, based on this survival curve, we concluded that the worsening stage in DS mice corresponds to the fourth week of life (P21 - P27), and that transition to the stabilization stage occurs during the fifth week. Moreover, the low frequency of spontaneous seizures prior to P19, the paucity of mortality before P20, and the susceptibility to thermal induction of seizures at P14 - P16 (Almog et al., 2019; Hawkins et al., 2017) suggest that the febrile stage corresponds to the third week of life (P14 - P20). Importantly, within this week, the epileptic phenotypes are gradually aggravated, beginning with susceptibility to febrile seizure (P14 - P16), progressing to infrequent spontaneous epileptic activity (P16 - P18), and ending with exacerbation to low incidence of premature mortality (P18 - P20). Despite that, the first half of this week (P14 - P18) still features mild epileptic phenotypes, characteristic of the febrile stage, especially when compared with those presented during the worsening stage (Fig. 1A, B).

Video-EEG recordings, taking place during the 12h light cycle and lasting 3 - 8 hours, were used to examine the epileptic (Fig. 1) and background EEG activity (Figs. 2, 3 see below) over the time course of the disease. Animals were monitored at three time points which correspond to the three stages of Dravet: (*i*) P18, the end of the febrile stage; (*ii*) during the fourth week of life, at the worsening stage of Dravet, or (*iii*) during the stabilization stage (P32 - P49). Mice younger than P16, at the time of the surgery, did not fully recover and therefore could not be recorded and included in these experiments. Interictal spikes were seen in all DS mice (Fig. 1C - F), tending towards a higher spike frequency in the worsening stage (Fig. 1C). Spontaneous seizures were recorded in one P18 DS mouse (16.6 % of the recorded DS mice at this age, Fig. 1D), four DS mice during the worsening stage (30.8%, Fig. 1E) and one DS mouse at the stabilization stage (8.3 %). Together, these data indicate that DS mice recapitulate the three stages of Dravet epilepsy, with progression from the febrile to the worsening stages occurring toward the end of the third week of life, which is subsequently followed by a stabilization stage beginning at the fifth week.

### Reduced EEG power in DS during the worsening stage

Despite the severe epileptic phenotypes of Dravet patients, background EEG activity appears mostly normal during the febrile and the beginning of the worsening stages, and therefore is not used for Dravet diagnosis (Wirrell et al., 2017). Here, to further identify alterations in background oscillations, we quantitatively analyzed background EEG activity of DS and WT littermates of different ages. To that end, five to ten 30 s long segments of the wake state, without movement artifact or epileptic activity, but following a movement as determined by the video recording, were analyzed and averaged for each mouse. In agreement with the onset of frequent spontaneous seizures and mortality, power spectral density (PSD) profiles were similar between WT and DS at the end of the febrile stage (Fig. 2A, B). In marked contrast, the worsening stage evidenced a global reduction of over 50% in total power (WT: 2.23 ± 0.31 μV^2^/Hz, n=14; DS: 1.03 ± 0.14 μV^2^/Hz, n=13; *p*=0.001). This reduction was observed across multiple frequency bands, with the sole exception of the gamma band (Fig. 2A, C, F). Finally, at the stabilization stage, total power was still reduced, but to a lower extent (WT: 1.55 ± 0.22 μV^2^/Hz, n=12; DS: 1.03 ± 0.08 μV^2^/Hz, n=11; *p*=0.043), with significant reductions mostly observed in the alpha and beta frequency bands (Fig. 2A, D, G).

Clinical power spectral analysis is usually based on the relative contribution of each frequency band to the total signal, normalizing the absolute power and masking any differences in total power (Tonner and Bein, 2006). This approach is more suitable for a clinical setting, as it is less affected by amplification settings of different systems used. When analyzing this way, the homogenous reduction in PSD at the worsening stage, from 0.5 Hz up to 30 Hz, resulted in similar percentage contribution (Fig. 2I), resembling data from young Dravet patients (up to 2 years as tested by (Holmes et al., 2012)). However, since the absolute power in the gamma band was not reduced in DS mice (Fig. 2F), its apparent relative contribution was higher following normalization (Fig. 2I). Later, during the stabilization stage, the contribution of gamma was similar in WT and DS mice (Fig, 2J). Despite that, at the stabilization stage there was a small reduction in the percentage of alpha band activity (Fig. 2J). Indeed, similar reductions in alpha activity were noted before in analysis of EEG background activity of Dravet patients, and were suggested to correlate with reduced cognitive functions (Holmes et al., 2012). Together, analysis of background EEG further supported the partition of Dravet epilepsy into three stages, showing near-normal background activity at the end of the febrile stage, dramatic reduction in total power during the worsening stage, and partial rectification at the stabilization stage.

### The risk for premature death correlates with lower PSD

To monitor PSD changes during development, a subset of WT and DS mice was recorded twice, at time points corresponding to the worsening and stabilization stages (P21 and P35) (Fig. 3). WT mice demonstrated an average total power reduction of ∼40%, with an identical trend observed throughout the tested cohort (Fig. 3A, B). In contrast, while DS mice, on average, also demonstrated power reduction (∼ 30%, Fig. 3C, D), the power was reduced in some mice, and increased in other mice (Fig. 3D). However, as 60% of the mice die during the fourth week of life (Fig. 1A), such longitudinal analysis inherently disregards these mice that die prematurely. Moreover, as frequent seizures have been shown to increase the risk of sudden unexpected death in epilepsy (SUDEP) (Kalume et al., 2013; Teran et al., 2019), this analysis also intrinsically selects mice with milder epilepsy or better compensatory mechanisms, which enable survival to the stabilization stage. A comparison of the PSD between mice that died prematurely and those that survived to the stabilization stage demonstrated decreased power in mice which eventually died during the worsening stage (Fig. 3C - E). Interestingly, the lower power was observed in multiple frequency bands, with the exception of the gamma band (Supplementary Fig. 1). Moreover, the relative spectral profile of these mice also differed, with lower contribution of theta and alpha but higher percentage of gamma band activity (Fig. 3F). In conclusion, these data demonstrate reduction in average PSD during maturation of both WT and DS mice. Nevertheless, with reduced power being a common motif in WT mice, power changes in DS demonstrated more variability. Moreover, low power at the worsening stage, together with increased gamma contribution, correlated with increased risk for SUDEP.

### Behavioral deficits in DS mice during the febrile, worsening and stabilization stages

Early development in Dravet patients is considered normal (Wirrell et al., 2017). Indeed, Dravet-associated comorbidities, including developmental delay, attention and cognitive deficits and motor impairments, manifest following the onset of severe epilepsy during the worsening stage. This progression suggests that epilepsy in Dravet plays a major role in the emergence of additional comorbidities (Brunklaus et al., 2012). This is further supported by normal EEG oscillations at the end of the febrile stage and the emergence of alterations in background activity during the worsening stage (Fig. 2). In contrast, several clinical studies reported subtle infantile developmental changes during the febrile stage, that may be concealed during traditional assessments (Chieffo et al., 2011; Guzzetta, 2011). Moreover, others suggested that developmental delay in Dravet may be independent of the epilepsy, and caused directly by the underlying *SCN1A* mutation (Nabbout et al., 2013; Ouss et al., 2019). To examine the relationship between Dravet epilepsy and Dravet associated comorbidities, and identify the onset of Dravet comorbidities, we performed behavioral tests in WT and DS mice.

In adult DS mice, motor deficits, hyperactivity and cognitive impairments are well documented (Han et al., 2012; Ito et al., 2013; Ricobaraza et al., 2019; Williams et al., 2019). However, the onset of these behavioral comorbidities remains unknown. In order to uncover their chronological timeline, we tested these behavioral features at time points corresponding to the different stages of Dravet. First, we tested infantile developmental milestones, at P8 - P11, before the febrile stage. Importantly, WT and DS mice performed similarly in these tests, which included early motor movement in an open arena and negative geotaxis, indicating that infantile development remains unaffected (Fig. 4 A, B). Next, additional behavioral assays, at timepoints corresponding to the three stages of Dravet, were performed. On the rotarod, which is designed to measure balance, coordination and grip strength (Brooks and Dunnett, 2009), early febrile stage DS mice (P14 - P16), which did not have recurrent seizures, demonstrated significantly shorter riding time-compared with WT mice. Interestingly, at the worsening stage, the fall latency was similar between the groups. Nevertheless, at the stabilization stage, DS mice demonstrated again reduced latency similar to the febrile stage (Fig. 4C).

**Figure 4.**
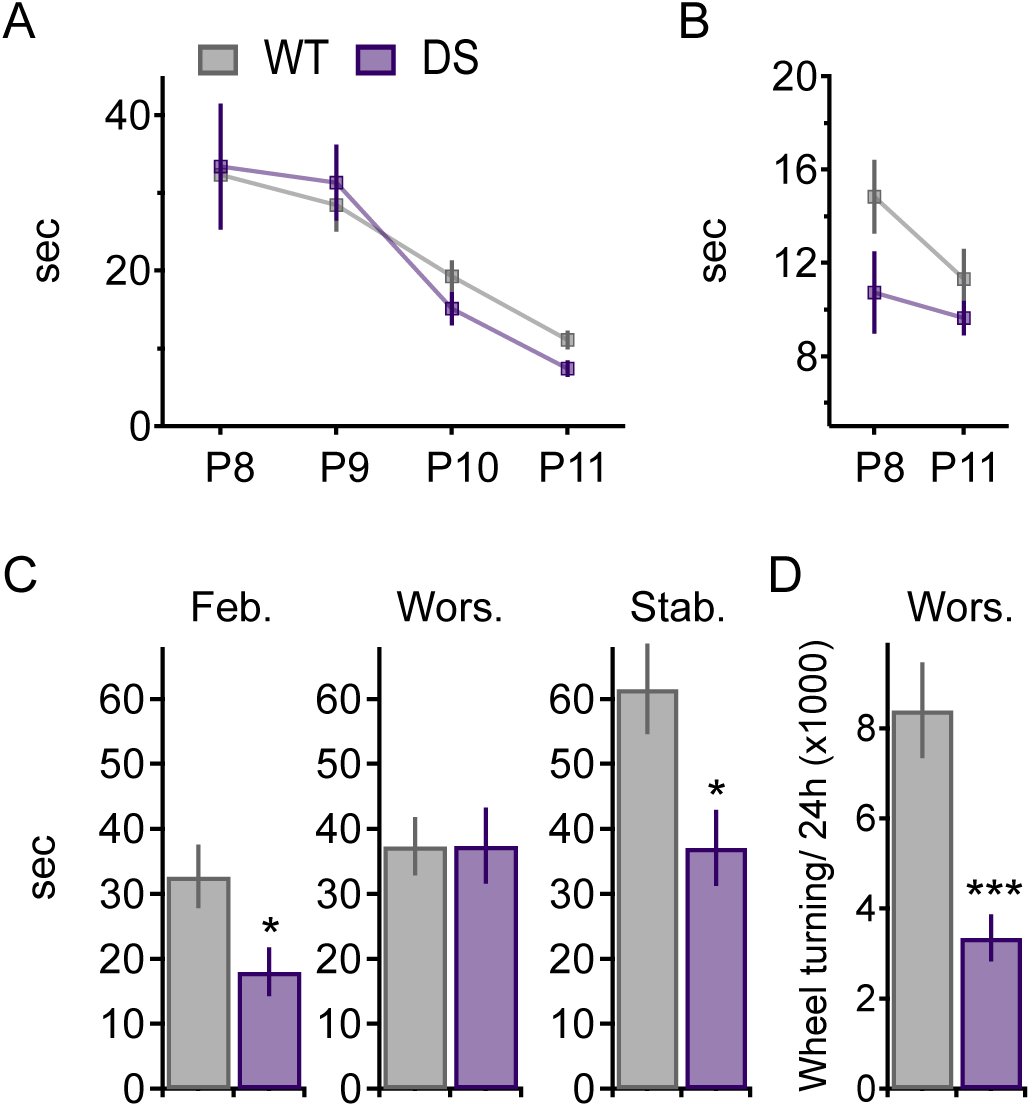
Motor impairment during the febrile stage. A - B, Early motor behaviors (P8 - P11) are similar between WT and DS mice. A, Mice, at the indicated ages, were placed at the center of a circular arena (13 cm diam.) and the time to exit the arena by crossing the border was recorded. WT: n = 9, DS: n = 9. B, Negative geotaxis at the indicated ages, the time to turn on inclined surface (45°) is depicted. WT: n = 8, DS: n = 10. C, Performance on the rotarod at the indicated timepoints. Febrile stage (P14 - P16), WT: n = 19, DS: n = 24; worsening stage (P21 - P27), WT: n = 25, DS: n = 15; stabilization stage (P35 - P40), WT: n = 18, DS: n = 9. See Supplementary Table 1 for separate averages of males and females. D, Voluntary running on home cage wheels during the worsening stage (P21 - P25), the average number of turns per day. WT: n = 8, DS: n = 8. \**p* < 0.05, \*\*\**p* < 0.001.

To further examine motor activity at the worsening stage, we monitored voluntary running behavior in the home cage (Fig. 4D). Intriguingly, despite the similar performance on the rotarod (Fig. 4C), DS mice had a fewer wheel turnings compared with WT (Fig. 4D), suggesting the existence of motor impairment also at the worsening stage, despite the slightly different presentation. Indeed, at the worsening stage, DS mice adopted a curious walking gait consisting of outwardly angled hind legs, which in turn may have helped their balance on the rotarod. Together, these data suggest that motor function impairments begin at the febrile stage, preceding the onset of severe epilepsy, and persisting to adulthood (Fig. 4C) (Ricobaraza et al., 2019). Thus, this comorbidity is likely not caused by recurrent seizures.

Next, we examined the behavior of the mice in the open field test. At the febrile stage (P14 - P16), DS mice had increased motor activity and traveled longer distances, and at higher velocities (Fig. 5A, B). This increased locomotion persisted from the febrile stage through the worsening and stabilization stages (Fig. 5A, B). These findings support the conclusion that locomotor hyperactivity in Dravet, similarly to motor impairment, are dissociated from the epilepsy. Analysis of mouse location within the open field arena can also be used to assess levels of anxiety, with reduced time spent at the center of the arena indicating elevated anxiety. Here, while WT and DS mice spent similar amounts of time in the center of the chamber during the febrile stage, high levels of anxiety persisted in DS mice through the worsening and the stabilization stages, while the time spent in the center of the arena gradually increased in WT mice (Fig. 5C). While these data can indicate a correlation between seizures and increased anxiety, the ability to detect differences in the anxiety level at the febrile stage may be limited. Indeed, the confoundingly high level of anxiety in pre-weaned mice (P14 - P16), renders monitoring differences between healthy and DS mice challenging (Fig. 5C).

**Figure 5.**
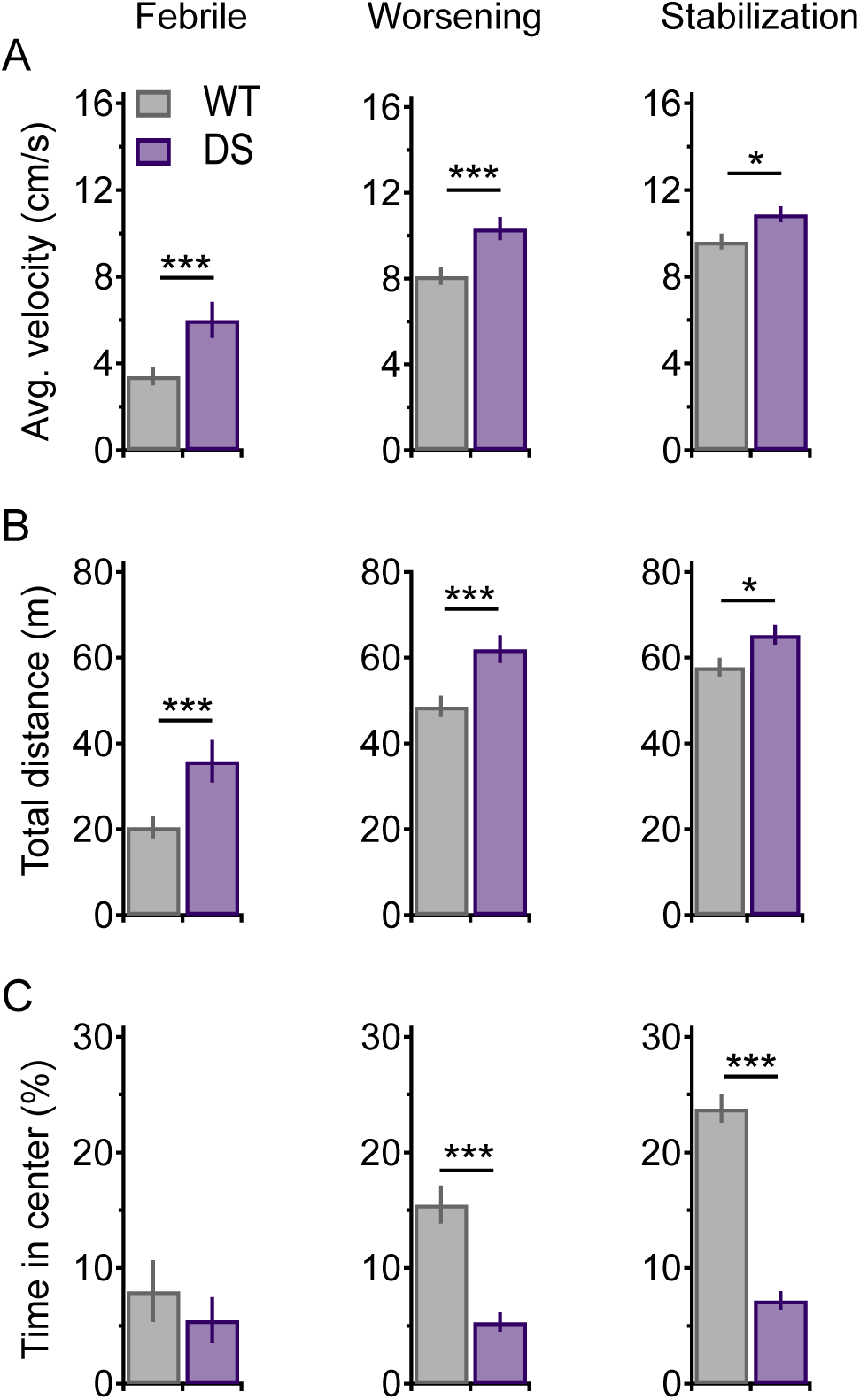
Hyperactivity in immature DS mice. A, Average velocity (cm/s) in open field at the indicated time points. B, Distance moved (m) in the open field. C, Time spent in the central portion of the arena. Febrile stage: P14 - P16: WT: n = 18, DS: n = 11; Worsening stage: P21 - P25: WT: n = 23, DS: n = 19, and stabilization stage: P35 - P49: WT: n = 38, DS: n = 30. See Supplementary Table 2 for separate averages of males and females. \**p* < 0.05, \*\**p* < 0.01, \*\*\**p* < 0.001.

Next, in order to explore the cognitive abilities at the different stages of Dravet, we performed the Y maze spontaneous alternation test. By relying on the natural exploration curiosity of mice, this behavioral assay examines spatial learning and memory. Beginning with P14 - P16 mice, we found that in this age 46% of the DS mice (6 out of 13 tested) and 36% of the WT (9 out of 25) were reluctant to explore the maze. However, mice just a few days older (P17 - P18), at the end of the febrile stage, but prior to changes in background EEG activity (Fig. 2), were willing to ambulate through the different arms of the maze. Despite that, the level of spontaneous alternation at this age did not rise above the level of random exploration (50% - chance level (Coutellier et al., 2012)) (Fig. 6A). Thus, the Y maze test may not be suitable for assessing cognitive abilities in rodents prior to P20 (Albani et al., 2014). Nevertheless, DS mice covered significantly longer distances during exploration (Fig. 6B), further corroborating DS hyperactivity at the febrile stage (Fig. 5A, B). Conversely, WT mice at the worsening stage displayed a slight, but significant, alternation level above chance level (52.7 % ± 1.15, One-Sample t-test relative to 50%, *p* = 0.024). Importantly, the level of alternation of DS mice at the same age was not above chance (48.2 % ± 1.5, One-Sample t-test, *p* = 0.23) (Fig. 6A), indicating impairment in cognitive abilities at the onset of epilepsy. At the stabilization stage, WT mice performed better, with a higher level of spontaneous alternation (56.8% ± 1.8, One-Sample t-test, *p* = 0.001). In contrast, alternation in DS mice still did not exceed chance level (46 % ± 3.4, One-Sample t-test, *p* = 0.25) (Fig. 6A).

**Figure 6.**
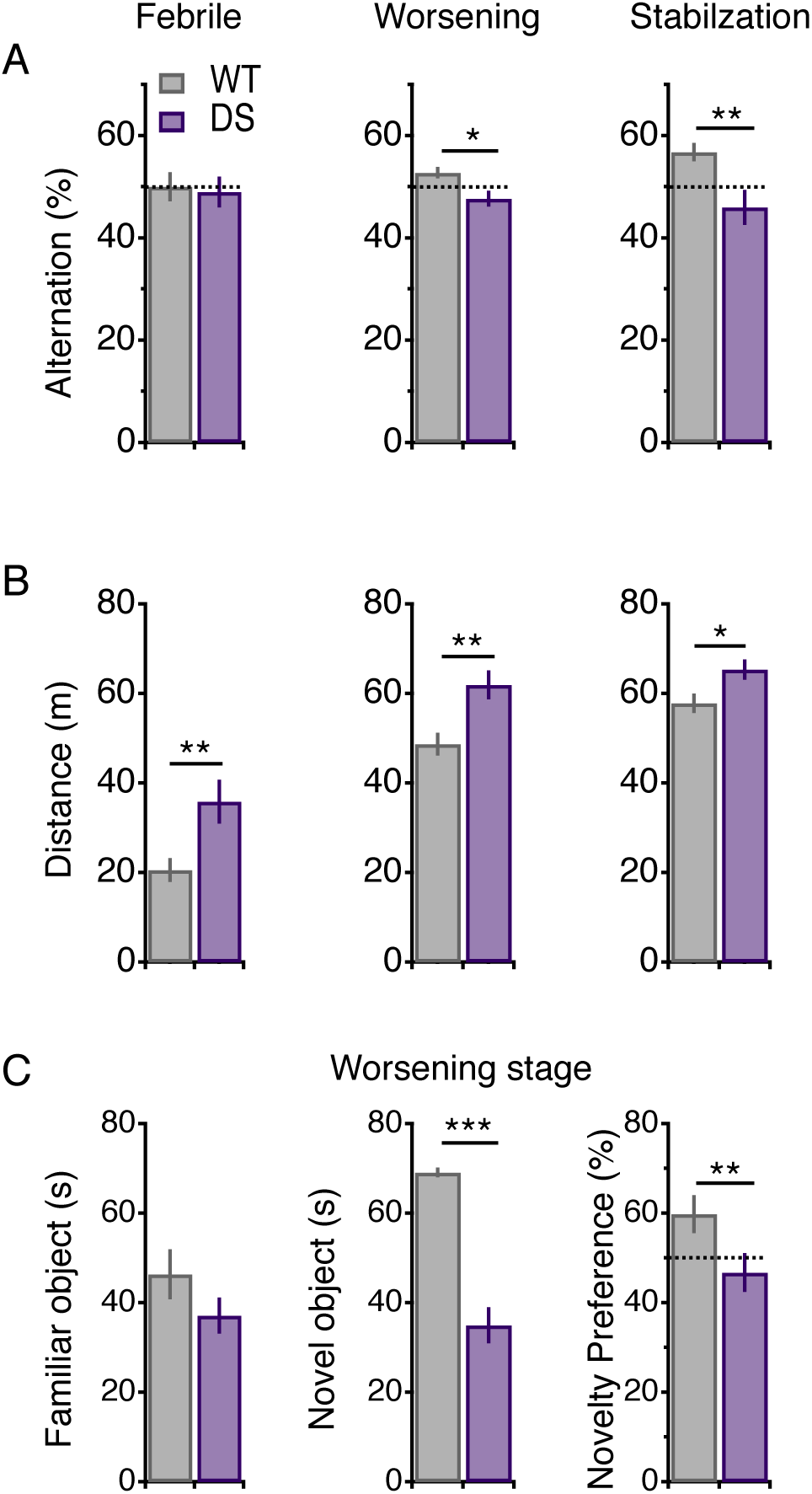
Reduced spontaneous alternations and preference for novelty coincide with the onset of epilepsy. A, Percent spontaneous alternation in the Y maze at the indicated timepoints, B, Average distance traveled. Febrile stage (P17 - P18). WT: n = 22, DS: n =11; Worsening stage (P21 - P25) WT: n = 30, DS: n =39, and stabilization stage (P35 - P49) WT: n = 23, DS: n = 16. See Supplementary Table 3 for separate averages of males and females. C, Performance in the novel object recognition test: time spent with either the familiar object (left) or novel object (middle), and novelty preference (right) are depicted. Worsening stage (P21 - P24) WT: n = 13, DS: n =13. \**p* < 0.05, \*\**p* < 0.01.

The low level of spontaneous alternation in WT mice, at the time point corresponding to the worsening stage (Fig. 6A), prompted us to further assess working memory at this age, using the novel object recognition test. In this test, while WT mice preferred exploring the novel object (novelty preference of 59.8 % ± 4.2, One-Sample t-test relative to 50%, *p* = 0.012), DS mice explored both objects similarly (46.8 % ± 4.3, One-Sample t-test, *p* = 0.14). These data, similarly to the results of the Y maze test, indicate that DS mice have cognitive deficits at the onset of severe epilepsy (Fig. 6C).

Together, these results indicate that reduced motor coordination and hyperactivity in the open field precede the onset of spontaneous seizures. This further indicates that some Dravet-associated comorbidities are not caused by the severe epilepsy. These early behavioral alterations are therefore likely to be related to neuronal changes that are driven by *Scn1a* mutation, independently from the epilepsy.

## Discussion

### Reduced EEG power at the worsening stage of Dravet

Seizure burden in Dravet changes through development. The febrile stage is characterized by normal development, albeit with febrile seizures and infrequent afebrile seizures. The disease then progresses to the worsening stage, with recurrent severe intractable seizures and developmental delay. In patients’ early school years, while the epilepsy improves, with reduced seizure burden, the other Dravet-associated comorbidities persist (Gataullina and Dulac, 2017).

Using DS mice, we demonstrate the full recapitulation of the developmental trajectory observed in patients. At the beginning of the third week of life (P14 - P16), DS mice are susceptible to thermal induction of seizures (Almog et al., 2019; Hawkins et al., 2017), however spontaneous seizures or premature mortality are rare (Fig. 1A) (Kang et al., 2018; Miller et al., 2014; Mistry et al., 2014; Ogiwara et al., 2013, 2007; Ricobaraza et al., 2019; Tsai et al., 2015; Williams et al., 2019; Yu et al., 2006). Transition into the worsening stage, after P18, is accompanied by the emergence of occasional spontaneous seizures (Fig. 1C, D). Nevertheless, background EEG profile at this age is similar to that of WT mice (Fig. 2), indicating that the epilepsy is still relatively mild at P18, just before the beginning of the worsening stage.

The worsening stage in DS mice corresponds to the fourth week of life (P21 - P27). It is characterized by frequent seizures and high premature mortality (Fig. 1). Importantly, other DS mouse models, based on global *Scn1a* truncation mutations, share an identical epileptic progression time course and overall mortality as the *Scn1a*^A1783V^ model used here (Kang et al., 2018; Mistry et al., 2014; Ogiwara et al., 2007; Tsai et al., 2015; Williams et al., 2019; Yu et al., 2006).

At the worsening stage, background EEG signals show a dramatic reduction in PSD, in multiple frequency bands (Fig. 2). Thus, as the total power correlated with the severity of epileptic phenotype and the risk for SUDEP (Fig. 3), these findings suggest that EEG power may serve as a marker for disease severity. In support of this, reduced EEG power has also been found to correlate with the severity of phenotype in Rett syndrome (Roche et al., 2019). Furthermore, in neonatal seizures and hypoxic ischemic encephalopathy, epileptic episodes during suppressed background EEG were correlated with more severe epilepsy (Korotchikova et al., 2011; Rowe et al., 1985).

During normal development, EEG power densities are increasing from infancy to early childhood, followed by a subsequent reduction towards sexual maturation (Aurlien et al., 2004; Bell et al., 2017). Importantly, we have also observed such a developmental trajectory in WT mice (Fig. 2). In contrast, overall developmental changes in power densities of DS mice were, on average, more stable (Fig. 2). Thus, these pronounced developmental differences may provide an *a priori* physical evidence for Dravet-associated developmental delay.

A longitudinal examination of individual DS mice, by recording their EEG activity at the worsening and stabilization stages, revealed divergent changes, with increased PSD in some mice but reduced PSD in others (Fig. 3D). On average, in a large cohort of mice, we found a small reduction in DS total power at the stabilization stage, compared to WT mice (Fig. 2D; of note, this small reduction was statistically significant only in the large cohort presented in Fig 2, but not in the smaller one presented in Fig. 3C, D). Interestingly, in support of our findings, reduced PSD was also demonstrated during non-rapid eye movement (NREM) sleep at the beginning of the stabilization stage (P30 - P35) (Kalume et al., 2015). Thus, reduced power in DS may be evident during both wakefulness and NREM sleep.

EEG power is linked to the balance between excitation and inhibition (Gao et al., 2017; Voytek and Knight, 2015; Womelsdorf et al., 2014). Hence, changes in PSD over the course of Dravet may be linked to the level of imbalance between these opposing neuronal activities. Indeed, previous electrophysiological studies suggested developmental changes in the function of inhibitory neurons (Almog et al., 2019; Favero et al., 2018; Mistry et al., 2014; Tsai et al., 2015). During the febrile stage, electrophysiological recordings in brain slices revealed a mild reduction in inhibition (Almog et al., 2019; De Stasi et al., 2016; Favero et al., 2018)(Tsai et al., 2015), in agreement with lack of change observed in background EEG (Fig. 2) or cortical LFP (De Stasi et al., 2016). In contrast, the worsening stage is characterized by a drastic reduction in the activity of inhibitory neurons (Almog et al., 2019; Han et al., 2012; Ogiwara et al., 2007; Rubinstein et al., 2015a, 2015b; Tsai et al., 2015; Yu et al., 2006), which may also manifest as the reduced EEG power observed (Fig. 2C, F). Finally, at the stabilization stage, the function of inhibitory neurons is partially restored (Almog et al., 2019; Favero et al., 2018), aligning well with the smaller differences in EEG PSD (Fig. 2D, G). Nevertheless, despite the lack of background activity alterations by the end of the febrile stage, DS mice still show susceptibility to thermally induced seizures (Almog et al., 2019; Hawkins et al., 2017), some epileptic activity (Fig. 1), and behavioral differences (motor impairment and hyperactivity in the open field, Figs. 4, 5, see below). Therefore, the lack of alterations in background EEG activity at this stagelikely indicates more subtle disturbances of excitation-inhibition balance, rather than lack of neuronal and network activity changes.

Previous studies demonstrated that increased seizure burden during the worsening stage correlates with increased risk for SUDEP (Kalume et al., 2013; Teran et al., 2019). Here, we add that reduced PSD also correlates with increased risk for SUDEP (Fig. 3). Together, these observations suggest that while the *Scn1a*^A1783V^ mutation is sufficient to cause Dravet, and all the mice develop epilepsy, some will show a higher seizure burden and greater risk for SUDEP. The neuronal basis for such disease severity variation, among mice that share the same *Scn1a* mutations and the same genetic background, is yet to be resolved. One possibility is that early changes, occurring already at the febrile stage, drive epileptogenic processes with some degree of flexibility inherent to this process.

Normalized PSD analysis of the spectral contribution demonstrated an increase in the percentage of the gamma band at the worsening stage (Fig. 2I, Fig. 3F). Importantly, the increased contribution of gamma also correlated with increased risk for SUDEP (Fig. 3F). This suggests that, in addition to low PSD, increased gamma may also be used as a marker for the severity of epilepsy. Corroborating this hypothesis, gamma band activity in epilepsy patients was shown to be higher in epileptic foci (Ren et al., 2015), and surgical removal of areas with high gamma resulted in a better outcome (Zweiphenning et al., 2019).

Gamma oscillations (30 - 100 Hz) are generated by a feed-forward inhibition activity, via (*i*) parvalbumin positive interneurons, (*ii*) disynaptic inhibition provided by Martinotti somatostatin positive cells, as well as (*iii*) thalamocortical connections (Womelsdorf et al., 2014). As reduction in the activity of both parvalbumin and somatostatin interneurons has been demonstrated in DS mice (De Stasi et al., 2016; Favero et al., 2018; Ogiwara et al., 2007; Rubinstein et al., 2015a; Tai et al., 2014), the preserved gamma power (leading to increased gamma contribution following normalization) is somewhat surprising. However, in contrast to the reduced function of cortical inhibitory neurons, thalamic reticular neurons were shown to be hyperactive in DS mice, leading to augmented thalamocortical oscillations (Ritter-Makinson et al., 2019). Thus, these opposing neuronal changes may be contributing to an overall gamma increase. Furthermore, increased gamma band contribution was also observed in stargazer mice (Maheshwari et al., 2016) and GAERS rats (Jones et al., 2010), animal models of absence epilepsy known to involve reduced function of inhibitory neurons (Maheshwari et al., 2013). Finally, computational models indicated that while reduced inhibition leads to decreased EEG power, gamma band activity remains unaffected, suggesting that the lower frequencies more sensitive to the balance between excitation and inhibition (Gao et al., 2017).

### Behavioral alterations in DS mice during the febrile stage

Early development of Dravet patients is considered normal, with clear signs of developmental delay appearing only after the onset of recurrent spontaneous seizures at the worsening stage (Gataullina and Dulac, 2017; Wirrell et al., 2017). Moreover, during the febrile stage, background EEG signals in both patients (Akiyama et al., 2010; Dravet et al., 2011; Genton et al., 2011; Holmes et al., 2012; Korff et al., 2007; Specchio et al., 2012; Takayama et al., 2014) and mice (Fig. 2) are also normal. Based on these observations, it was suggested that the severe frequent seizures contribute to cognitive decline and the other Dravet-associated comorbidities (Ben-Ari and Holmes, 2006; Cetica et al., 2017). Other clinical studies, however, have suggested that developmental delay is epilepsy-independent, indicating that cognitive deficits cannot be prevented, even with adequate seizure control (Brunklaus and Zuberi, 2014; Nabbout et al., 2013; Ouss et al., 2019).

Here, we performed behavioral tests in DS and WT littermates in order to trace the onset of motor impairment, hyperactivity and cognitive deficits, and any temporal correlation to epilepsy. While DS mice demonstrated typical early developmental milestones (Fig. 4A, B), behavioral deficits were presented at the febrile stage, prior to the onset of severe spontaneous seizures (Figs. 4 - 6). These included reduced performance on the rotarod, as well as hyperactivity in the open field and the Y maze (Figs. 4 - 6). These results suggest that these behavioral phenotypes are independent of epilepsy. Moreover, the rotarod and open field tests were performed at P14 - P16, the very beginning of the febrile stage, prior to the appearance of spontaneous seizures or premature death (Fig. 1A and (Kang et al., 2018; Mistry et al., 2014; Ogiwara et al., 2007; Tsai et al., 2015; Williams et al., 2019; Yu et al., 2006). Importantly, previous measurements of the levels of Na_V_1.1, encoded by the *SCN1A* gene, revealed low expression early in life, which is later increasing, coinciding with the onset of the febrile stage of Dravet (Cheah et al., 2013). Thus, we propose that motor impairment and hyperactivity in the open field, as well as the mild epileptic phenotypes during the febrile stage, are all, independently, caused by altered neuronal and network activity due to Na_V_1.1 loss of function. These findings also support the notion that early development in Dravet, during the febrile stage, conceals quiescent behavioral alterations. In corroboration of this, early visuomotor dysfunctions were noted in patients even before the onset of epilepsy (Battaglia et al., 2016; Chieffo et al., 2011; Guzzetta, 2011; Ricci et al., 2015; Verheyen et al., 2019).

Interestingly, increased anxiety and deficits in working memory were not detected before the worsening stage, coinciding with the onset of severe seizures (Figs. 5, 6). These deficits persisted into the stabilization stage (Figs. 5, 6) and adulthood (Han et al., 2012; Ito et al., 2013; Ricobaraza et al., 2019). Unfortunately, assessing the levels of anxiety and working memory at the febrile stage may be confounded by the behavioral features of immature mice and the limited sensitivity of these behavioral tests (Figs. 5, 6). Nevertheless, studies of adult DS mice, featuring selective deletion of *Scn1a* in parvalbumin- and somatostatin-expressing inhibitory neurons, reported mild epilepsy with cognitive deficit (Rubinstein et al., 2015a; Tatsukawa et al., 2018). Similarly, local deletions of *Scn1a* in the hippocampus caused either mild or no epilepsy, with cognitive deficits (Bender et al., 2016, 2013; Stein et al., 2019). These results suggest that cognitive deficits, similarly to motor impairment and hyperactivity may also be independent of the epilepsy. Together, these data indicate that *Scn1a* mutations are sufficient to cause at least some of Dravet-associated comorbidities, regardless of additional ramifications that may result from recurrent seizures.

## Acknowledgments

We wish to thank Dr. Yoni Haitin and Dr. Moshe Giladi for helpful discussions and critical reading of the manuscript. We also wish to thank Dr. Lior Bikovski and the Myers Neuro-Behavioral Core Facility at Tel Aviv University. This work was supported by the Israel Science Foundation (grant 1454/17; M.R), The Fritz Thyssen Stiftung (grant 10.17.1.023MN; M.R), The Jérôme Lejeune Foundation (grant #1565 - 2016b; M.R), The V. Schreiber Research Fund and The Aya Baharav Fund of the Sackler School of Medicine, Tel Aviv University (M.R).

## References

Akiyama, M., Kobayashi, K., Yoshinaga, H., Ohtsuka, Y., 2010. A long-term follow-up study of Dravet syndrome up to adulthood. Epilepsia 51, 1043–1052.

Albani, S.H., McHail, D.G., Dumas, T.C., 2014. Developmental studies of the hippocampus and hippocampal-dependent behaviors: Insights from interdisciplinary studies and tips for new investigators. Neurosci. Biobehav. Rev.

Almog, Y., Brusel, M., Anderson, K., Rubinstein, M., 2019. Early hippocampal hyperexcitability followed by disinhibition in a mouse model of Dravet syndrome. bioRxiv 790170.

Aurlien, H., Gjerde, I.O., Aarseth, J.H., Eldøen, G., Karlsen, B., Skeidsvoll, H., Gilhus, N.E., 2004. EEG background activity described by a large computerized database. Clin. Neurophysiol. 115, 665–673.

Battaglia, D., Ricci, D., Chieffo, D., Guzzetta, F., 2016. Outlining a core neuropsychological phenotype for Dravet syndrome. Epilepsy Res. 120, 91–97.

Bell, M.A., Wolfe, C.D., Wolfe, C.D., 2017. Changes in brain functioning from infancy to early childhood: Evidence from EEG power and coherence during working memory tasks. Dev. Neuropsychol. 5641, 20–38.

Ben-Ari, Y., Holmes, G.L., 2006. Effects of seizures on developmental processes in the immature brain. Lancet Neurol. 5, 1055–1063.

Bender, A.C., Luikart, B.W., Lenck-Santini, P.P., 2016. Cognitive deficits associated with Nav1.1 alterations: Involvement of neuronal firing dynamics and oscillations. PLoS One 11, e0151538.

Bender, A.C., Natola, H., Ndong, C., Holmes, G.L., Scott, R.C., Lenck-Santini, P.P., 2013. Focal *Scn1a* knockdown induces cognitive impairment without seizures. Neurobiol. Dis. 54, 297– 307.

Brooks, S.P., Dunnett, S.B., 2009. Tests to assess motor phenotype in mice: A user’s guide. Nat. Rev. Neurosci. 10, 519–529.

Brunklaus, A., Ellis, R., Reavey, E., Forbes, G.H., Zuberi, S.M., 2012. Prognostic, clinical and demographic features in *SCN1A* mutation-positive Dravet syndrome. Brain 135, 2329–2336.

Brunklaus, A., Zuberi, S.M., 2014. Dravet syndrome-From epileptic encephalopathy to channelopathy. Epilepsia 55, 979–984.

Cetica, V., Chiari, S., Mei, D., Parrini, E., Grisotto, L., Marini, C., Pucatti, D., Ferrari, A., Sicca, F., Specchio, N., Trivisano, M., Battaglia, D., Contaldo, I., Zamponi, N., Petrelli, C., Granata, T., Ragona, F., Avanzini, G., Guerrini, R., 2017. Clinical and genetic factors predicting Dravet syndrome in infants with *SCN1A* mutations. Neurology 88, 1037–1044.

Cheah, C.S., Westenbroek, R.E., Roden, W.H., Kalume, F., Oakley, J.C., Jansen, L.A., Catterall, W.A., 2013. Correlations in timing of sodium channel expression, epilepsy, and sudden death in Dravet syndrome. Channels. 7, 468–472.

Cheah, C.S., Yu, F.H., Westenbroek, R.E., Kalume, F.K., Oakley, J.C., Potter, G.B., Rubenstein, J.L., Catterall, W.A., 2012. Specific deletion of Na_V_1.1 sodium channels in inhibitory interneurons causes seizures and premature death in a mouse model of Dravet syndrome. Proc. Natl. Acad. Sci. 109, 14646–14651.

Chieffo, D., Ricci, D., Baranello, G., Martinelli, D., Veredice, C., Lettori, D., Battaglia, D., Dravet, C., Mercuri, E., Guzzetta, F., 2011. Early development in Dravet syndrome; visual function impairment precedes cognitive decline. Epilepsy Res. 93, 73–79.

Claes, L., Del-Favero, J., Ceulemans, B., Lagae, L., Van Broeckhoven, C., De Jonghe, P., 2002. De novo mutations in the sodium-channel gene *SCN1A* cause severe myoclonic epilepsy of infancy. Am. J. Hum. Genet. 68, 1327–1332.

Coutellier, L., Beraki, S., Ardestani, P.M., Saw, N.L., Shamloo, M., 2012. Npas4: A Neuronal transcription factor with a key role in social and cognitive functions relevant to developmental disorders. PLoS One 7. e46604

De Stasi, A.M., Farisello, P., Marcon, I., Cavallari, S., Forli, A., Vecchia, D., Losi, G., Mantegazza, M., Panzeri, S., Carmignoto, G., Bacci, A., Fellin, T., 2016. Unaltered Network Activity and Interneuronal Firing during Spontaneous Cortical Dynamics in Vivo in a Mouse Model of Severe Myoclonic Epilepsy of Infancy. Cereb. Cortex 26, 1778–1794.

Dravet, C., Bureau, M., Bernardina, B.D., Guerrini, R., 2011. Severe myoclonic epilepsy in infancy (Dravet syndrome) 30 years later. Epilepsia 52, 1–2.

Dravet, C., Oguni, H., 2013. Dravet syndrome (severe myoclonic epilepsy in infancy). Handb. Clin. Neurol. 111, 627–633.

Favero, M., Sotuyo, N.P., Lopez, E., Kearney, J.A., Goldberg, E.M., 2018. A transient developmental window of fast-spiking interneuron dysfunction in a mouse model of Dravet syndrome. J. Neurosci. 38, 7912–7927.

Gao, R., Peterson, E.J., Voytek, B., 2017. Inferring synaptic excitation/inhibition balance from field potentials. Neuroimage 158, 70–78.

Gataullina, S., Dulac, O., 2017. From genotype to phenotype in Dravet disease. Seizure. 44, 58– 64

Genton, P., Velizarova, R., Dravet, C., 2011. Dravet syndrome: the long-term outcome. Epilepsia 52, 44–49.

Guzzetta, F., 2011. Cognitive and behavioral characteristics of children with Dravet syndrome: an overview. Epilepsia 52 Suppl 2, 35.

Han, S., Tai, C., Westenbroek, R.E., Yu, F.H., Cheah, C.S., Potter, G.B., Rubenstein, J.L., Scheuer, T., De La Iglesia, H.O., Catterall, W.A., 2012. Autistic-like behaviour in *Scn1a* ^+/-^ mice and rescue by enhanced GABA-mediated neurotransmission. Nature 489, 385–390.

Hawkins, N.A., Anderson, L.L., Gertler, T.S., Laux, L., George, A.L., Kearney, J.A., 2017. Screening of conventional anticonvulsants in a genetic mouse model of epilepsy. Ann. Clin. Transl. Neurol. 4, 326–339.

Holmes, G.L., Bender, A.C., Wu, E.X., Scott, R.C., Lenck-Santini, P.P., Morse, R.P., 2012. Maturation of EEG oscillations in children with sodium channel mutations. Brain Dev. 34, 469–477.

Ito, S., Ogiwara, I., Yamada, K., Miyamoto, H., Hensch, T.K., Osawa, M., Yamakawa, K., 2013. Mouse with Na v 1.1 haploinsufficiency, a model for Dravet syndrome, exhibits lowered sociability and learning impairment. Neurobiol. Dis. 49, 29–40.

Jones, N.C., Martin, S., Megatia, I., Hakami, T., Salzberg, M.R., Pinault, D., Morris, M.J., O’Brien, T.J., van den Buuse, M., 2010. A genetic epilepsy rat model displays endophenotypes of psychosis. Neurobiol. Dis. 39, 116–125.

Kalume, F., Oakley, J.C., Westenbroek, R.E., Gile, J., de la Iglesia, H.O., Scheuer, T., Catterall, W.A., 2015. Sleep impairment and reduced interneuron excitability in a mouse model of Dravet Syndrome. Neurobiol. Dis. 77, 141–154.

Kalume, F., Westenbroek, R.E., Cheah, C.S., Yu, F.H., Oakley, J.C., Scheuer, T., Catterall, W.A., 2013. Sudden unexpected death in a mouse model of Dravet syndrome. J. Clin. Invest. 123,

Kalume, F., Yu, F.H., Westenbroek, R.E., Scheuer, T., Catterall, W.A., 2007. Reduced sodium current in Purkinje neurons from NaV1.1 mutant mice: Implications for ataxia in severe myoclonic epilepsy in infancy. J. Neurosci. 27, 11065–11074.

Kang, S.K., Hawkins, N.A., Kearney, J.A., 2018. C57BL/6J and C57BL/6N substrains differentially influence phenotype severity in the *Scn1a*^+/−^ mouse model of Dravet syndrome. Epilepsia Open 4, epi4.12287.

Korff, C., Laux, L., Kelley, K., Goldstein, J., Koh, S., Nordli, D., Christian, K., Linda, L., Kent, K., Joshua, G., Sookyong, K., Douglas Jr, N., 2007. Dravet syndrome (severe myoclonic epilepsy in infancy): a retrospective study of 16 patients. J. Child Neurol. 22, 185–194.

Korotchikova, I., Stevenson, N.J., Walsh, B.H., Murray, D.M., Boylan, G.B., 2011. Quantitative EEG analysis in neonatal hypoxic ischaemic encephalopathy. Clin. Neurophysiol. 122, 1671–1678.

Kuo, F.-S., Cleary, C.M., LoTurco, J.J.L., Chen, X., Mulkey, D.K., 2019. Disordered breathing in a mouse model of Dravet syndrome. Elife 8, e43387.

Maheshwari, A., Marks, R.L., Yu, K.M., Noebels, J.L., 2016. Shift in interictal relative gamma power as a novel biomarker for drug response in two mouse models of absence epilepsy. Epilepsia 57, 79–88.

Maheshwari, A., Nahm, W.K., Noebels, J.L., 2013. Paradoxical proepileptic response to NMDA receptor blockade linked to cortical interneuron defect in stargazer mice. Front. Cell. Neurosci. 7. doi: 10.3389/fncel.2013.00156.

Miller, A.R., Hawkins, N.A., McCollom, C.E., Kearney, J.A., 2014. Mapping genetic modifiers of survival in a mouse model of Dravet syndrome. Genes, Brain Behav. 13, 163–172.

Mistry, A.M., Thompson, C.H., Miller, A.R., Vanoye, C.G., George, A.L., Kearney, J.A., 2014. Strain- and age-dependent hippocampal neuron sodium currents correlate with epilepsy severity in Dravet syndrome mice. Neurobiol. Dis. 65, 1–11.

Nabbout, R., Chemaly, N., Chipaux, M., Barcia, G., Bouis, C., Dubouch, C., Leunen, D., Jambaqué, I., Dulac, O., Dellatolas, G., Chiron, C., 2013. Encephalopathy in children with Dravet syndrome is not a pure consequence of epilepsy. Orphanet J. Rare Dis. 8, 176.

Ogiwara, I., Iwasato, T., Miyamoto, H., Iwata, R., Yamagata, T., Mazaki, E., Yanagawa, Y., Tamamaki, N., Hensch, T.K., Itohara, S., Yamakawa, K., 2013. Nav1.1 haploinsufficiency in excitatory neurons ameliorates seizure-associated sudden death in a mouse model of dravet syndrome. Hum. Mol. Genet. 22, 4784–4804.

Ogiwara, I., Miyamoto, H., Morita, N., Atapour, N., Mazaki, E., Inoue, I., Takeuchi, T., Itohara, S., Yanagawa, Y., Obata, K., Furuichi, T., Hensch, T.K., Yamakawa, K., 2007. Nav1.1 localizes to axons of parvalbumin-positive inhibitory interneurons: a circuit basis for epileptic seizures in mice carrying an Scn1a gene mutation. J. Neurosci. 27, 5903–5914.

Ouss, L., Leunen, D., Laschet, J., Chemaly, N., Barcia, G., Losito, E.M., Aouidad, A., Barrault, Z., Desguerre, I., Breuillard, D., Nabbout, R., 2019. Autism spectrum disorder and cognitive profile in children with Dravet syndrome: Delineation of a specific phenotype. Epilepsia Open 4, 40–53.

Ren, L., Kucewicz, M.T., Cimbalnik, J., Matsumoto, J.Y., Brinkmann, B.H., Hu, W., Marsh, W.R., Meyer, F.B., Stead, S.M., Worrell, G.A., 2015. Gamma oscillations precede interictal epileptiform spikes in the seizure onset zone. Neurology 84, 602–608.

Ricci, D., Chieffo, D., Battaglia, D., Brogna, C., Contaldo, I., De Clemente, V., Losito, E., Dravet, C., Mercuri, E., Guzzetta, F., 2015. A prospective longitudinal study on visuo-cognitive development in Dravet syndrome: Is there a “dorsal stream vulnerability”? Epilepsy Res. 109, 57–64.

Ricobaraza, A., Mora-Jimenez, L., Puerta, E., Sanchez-Carpintero, R., Mingorance, A., Artieda, J., Nicolas, M.J., Besne, G., Bunuales, M., Gonzalez-Aparicio, M., Sola-Sevilla, N., Valencia, M., Hernandez-Alcoceba, R., 2019. Epilepsy and neuropsychiatric comorbidities in mice carrying a recurrent Dravet syndrome *SCN1A* missense mutation. Sci. Rep. 9, 14172.

Ritter-Makinson, S., Clemente-Perez, A., Higashikubo, B., Cho, F.S., Holden, S.S., Bennett, E., Chkaidze, A., Eelkman Rooda, O.H.J., Cornet, M.C., Hoebeek, F.E., Yamakawa, K., Cilio, M.R., Delord, B., Paz, J.T., 2019. Augmented reticular thalamic bursting and seizures in *Scn1a*-Dravet syndrome. Cell Rep. 26, 54–64.e6.

Roche, K.J., Leblanc, J.J., Levin, A.R., O’Leary, H.M., Baczewski, L.M., Nelson, C.A., 2019. Electroencephalographic spectral power as a marker of cortical function and disease severity in girls with Rett syndrome. J. Neurodev. Disord. 11. doi:10.1186/s11689-019-9275-z

Rowe, J.C., Holmes, G.L., Hafford, J., Baboval, D., Robinson, S., Philipps, A., Rosenkrantz, T., Raye, J., 1985. Prognostic value of the electroencephalogram in term and preterm infants following neonatal seizures. Electroencephalogr. Clin. Neurophysiol. 60, 183–196.

Rubinstein, M., Han, S., Tai, C., Westenbroek, R.E., Hunker, A., Scheuer, T., Catterall, W.A., 2015a. Dissecting the phenotypes of Dravet syndrome by gene deletion. Brain 138, 2219– 2233.

Rubinstein, M., Westenbroek, R.E., Yu, F.H., Jones, C.J., Scheuer, T., Catterall, W.A., 2015b. Genetic background modulates impaired excitability of inhibitory neurons in a mouse model of Dravet syndrome. Neurobiol. Dis. 73, 106–117.

Schwenk, F., Baron, U., Rajewsky, K., 1995. A cre -transgenic mouse strain for the ubiquitous deletion of loxP -flanked gene segments including deletion in germ cells. Nucleic Acids Res. 23, 5080–5081.

Specchio, N., Balestri, M., Trivisano, M., Japaridze, N., Striano, P., Carotenuto, A., Cappelletti, S., Specchio, L.M., Fusco, L., Vigevano, F., 2012. Electroencephalographic features in Dravet syndrome: Five-year follow-up study in 22 patients. J. Child Neurol. 27, 439–444.

Stein, R.E., Kaplan, J.S., Li, J., Catterall, W.A., 2019. Hippocampal deletion of NaV 1.1 channels in mice causes thermal seizures and cognitive deficit characteristic of Dravet Syndrome. Proc. Natl. Acad. Sci. 116, 16571–16576

Styr, B., Gonen, N., Zarhin, D., Ruggiero, A., Atsmon, R., Gazit, N., Braun, G., Frere, S., Vertkin, I., Shapira, I., Harel, M., Heim, L.R., Katsenelson, M., Rechnitz, O., Fadila, S., Derdikman, D., Rubinstein, M., Geiger, T., Ruppin, E., Slutsky, I., 2019. Mitochondrial regulation of the hippocampal firing rate set point and seizure susceptibility. Neuron 102, 1009–1024.e8.

Tai, C., Abe, Y., Westenbroek, R.E., Scheuer, T., Catterall, W.A., 2014. Impaired excitability of somatostatin- and parvalbumin-expressing cortical interneurons in a mouse model of Dravet syndrome. Proc. Natl. Acad. Sci. 111, E3139–E3148.

Takayama, R., Fujiwara, T., Shigematsu, H., Imai, K., Takahashi, Y., Yamakawa, K., Inoue, Y., 2014. Long-term course of Dravet syndrome: A study from an epilepsy center in Japan. Epilepsia 55, 528–538.

Tatsukawa, T., Ogiwara, I., Mazaki, E., Shimohata, A., Yamakawa, K., 2018. Impairments in social novelty recognition and spatial memory in mice with conditional deletion of *Scn1a* in parvalbumin-expressing cells. Neurobiol. Dis. 112, 24–34.

Teran, F.A., Kim, Y., Crotts, M.S., Bravo, E., Emaus, K.J., Richerson, G.B., 2019. Time of Day and a Ketogenic Diet Influence Susceptibility to SUDEP in *Scn1a*^R1407X/+^ Mice. Front. Neurol. 10. https://doi.org/10.3389/fneur.2019.00278

Tonner, P.H., Bein, B., 2006. Classic electroencephalographic parameters: Median frequency, spectral edge frequency etc. Best Pract. Res. Clin. Anaesthesiol. 20, 147–159.

Tsai, M.S., Lee, M.L., Chang, C.Y., Fan, H.H., Yu, I.S., Chen, Y.T., You, J.Y., Chen, C.Y., Chang, F.C., Hsiao, J.H., Khorkova, O., Liou, H.H., Yanagawa, Y., Lee, L.J., Lin, S.W., 2015. Functional and structural deficits of the dentate gyrus network coincide with emerging spontaneous seizures in an *Scn1a* mutant dravet syndrome model during development. Neurobiol. Dis. 77, 35–48.

Verheyen, K., Verbecque, E., Ceulemans, B., Schoonjans, A., Van De Walle, P., Hallemans, A., 2019. Motor development in children with Dravet syndrome. Dev. Med. Child Neurol. 61, 950–956.

Voytek, B., Knight, R.T., 2015. Dynamic Network Communication as a Unifying Neural Basis for Cognition, Development, Aging, and Disease. Biol. Psychiatry 77, 1089–1097.

Williams, A.D., Kalume, F., Westenbroek, R.E., Catterall, W.A., 2019. A more efficient conditional mouse model of Dravet syndrome: Implications for epigenetic selection and sex-dependent behaviors. J. Neurosci. Methods 325, 108315.

Wirrell, E.C., Laux, L., Donner, E., Jette, N., Knupp, K., Meskis, M.A., Miller, I., Sullivan, J., Welborn, M., Berg, A.T., 2017. Optimizing the diagnosis and management of Dravet syndrome: recommendations from a north american consensus panel. Pediatr. Neurol. 68, 18–34.e3.

Womelsdorf, T., Valiante, T.A., Sahin, N.T., Miller, K.J., Tiesinga, P., 2014. Dynamic circuit motifs underlying rhythmic gain control, gating and integration. Nat. Neurosci. 17,1031– 1039.

Yu, F.H., Mantegazza, M., Westenbroek, R.E., Robbins, C.A., Kalume, F., Burton, K.A., Spain, W.J., McKnight, G.S., Scheuer, T., Catterall, W.A., 2006. Reduced sodium current in GABAergic interneurons in a mouse model of severe myoclonic epilepsy in infancy. Nat. Neurosci. 9, 1142–1149.

Zweiphenning, W.J.E.M., Keijzer, H.M., van Diessen, E., van ‘t Klooster, M.A., van Klink, N.E.C., Leijten, F.S.S., van Rijen, P.C., van Putten, M.J.A.M., Braun, K.P.J., Zijlmans, M., 2019. Increased gamma and decreased fast ripple connections of epileptic tissue: A high-frequency directed network approach. Epilepsia 60, 1908–1920.

