## Supplementary Fig. 1, Supplementary Table 1, Supplementary Table 2, Supplementary Table 3 for "Early EEG and behavioral alterations in Dravet mice"

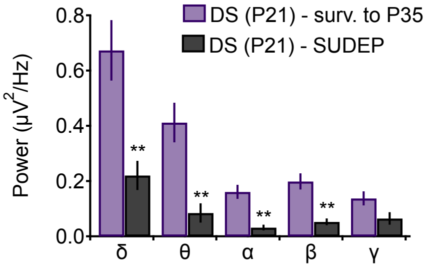


Supplementary Fig. 1. Related to Fig. 3. Total power in each frequency band during the worsening stage (δ: 0.5 - 4 Hz; θ: 4 - 8 Hz; α: 8 -12 Hz; β: 12 - 30 Hz; γ: 30 - 100 Hz) in DS mice that survived to the stabilization stage and DS mice that died from SUDEP during the worsening stage. $**p < 0.01$.

Supplementary Table 1. Rotarod - additional data related to Fig. 4

| | Latency to fall (s) | Latency to fall (s) | Latency to fall (s) |
|---|---|---|---|
| | P14-P16 | P21-P25 | P35-P49 |
| **WT male** | 31.21 ± 9.29 (n=8) | 26.48 ± 5.08 (n=11) | 42.87 ± 4.45 (n=8) |
| **WT female** | 35.08 ± 5.97 (n=10) | 45.93 ± 6.2 (n=14) | 82.03 ± 10.5 (n=9) |
| **DS male** | 21.49 ± 6.52 (n=13) | 26.13 ± 7.97 (n=5) | 36.01 ± 7.42 (n=5) |
| **DS female** | 14.99 ± 2.39 (n=13) | 43.15 ± 7.35 (n=10) | 36.6 ± 8.3 (n=5) |

Supplementary Table 2. Open field - additional data related to Fig. 5

| | Distance (m) | Velocity (cm/s) | Distance (m) | Velocity (cm/s) | Distance (m) | Velocity (cm/s) |
|---|---|---|---|---|---|---|
| | P14-P16 | | P21-P25 | | P35-P49 | |
| **WT male** | 17.7 ± 2.7 | 2.9 ± 0.4 (n=10) | 51.3 ± 4.6 | 8.5 ± 0.8 (n=10) | 57.3 ± 3.5 | 9.6 ± 0.6 (n=20) |
| **WT female** | 24.2 ± 4.7 | 4 ± 0.8 (n=8) | 47 ± 2.8 | 7.8 ± 0.5 (n=14) | 58.4 ± 2.7 | 9.7 ± 0.4 (n=19) |
| **DS male** | 40 ± 5.8 | 6.7 ± 1 (n=7) | 68 ± 4.5 | 11.4 ± 0.7 (n=11) | 66.1 ± 3.3 | 11 ± 0.6 (n=13) |
| **DS female** | 28 ± 8.6 | 4.7 ± 1.4 (n=4) | 53.4 ± 2.5 | 9 ± 0.4 (n=8) | 64.9 ± 3.1 | 10.8 ± 0.5 (n=17) |

Supplementary Table 3. Y maze spontaneous alternation - additional data related to Fig 6.

| | Distance (m) | Alternation (%) | Distance (m) | Alternation (%) | Distance (m) | Alternation (%) |
|---|---|---|---|---|---|---|
| | P17-P18 | | P21-P25 | | P35-P49 | |
| **WT male** | 25.5 ± 2.4 | 45.5 ± 3.4 (n=12) | 39.6 ± 2.2 | 50.8 ± 1.6 (n=16) | 37.7 ± 2.7 | 61.4 ± 2.3 (n=11) |
| **WT female** | 25.2± 1.9 | 55.4 ± 4.4 (n=10) | 36.6 ± 2 | 54.8 ± 1.5 (n=15) | 48.9± 2.9 | 52.1± 2.6 (n=10) |
| **DS male** | 41.5 ± 1.2 | 45.4 ± 3.4 (n=8) | 52.4 ± 2.8 | 44.7 ± 3 (n=19) | 51.5 ± 3.7 | 43.8 ± 3.9 (n=10) |
| **DS female** | 57.1 ± 8.7 | 55.7 ± 5.7 (n=5) | 45.2 ± 2.7 | 49.5 ± 2.1 (n=21) | 48.9 ± 3.1 | 49.7 ± 5.1 (n=6) |
